# Incorporating multi-scale module kernel for disease-gene identification in biological networks

**DOI:** 10.1101/2022.07.28.501869

**Authors:** Ju Xiang, Kaixin Zeng, Shengkai Chen, Xiangmao Meng, Ruiqing Zheng, Ying Zheng, Yahui Long, Min Li

## Abstract

Biomedical data mining plays a crucial role in studying diseases, with disease-gene identification being one of the most prominent areas of research in this field. Many biomolecule networks are known to have multi-scale module structures, which may be helpful for studying complex diseases, but the mining and utilization of multi-scale module structure is an open issue. Therefore, we present a kind of novel hybrid network-based method (HyMSMK) for disease-gene identification through incorporating multi-scale module kernel in biomolecule networks. We first apply exponential sampling to construct multi-scale module profile containing local to global structural information, where modules at different scales are extracted from comprehensive interactome by multi-scale modularity optimization. Then, the multi-scale module profile is preprocessed by the relative information content, and is used to generate multi-scale module kernel, which is further preprocessed by kernel sparsification. We design multiple schemes for incorporating multi-scale module kernel to discover potential disease-related genes. We investigate the performance of these schemes by experimental evaluations, show the positive effect of kernel sparsification on reducing the requirement for space and time, and confirm the superior performance of our method compared to other state-of-art network-based baselines. The study demonstrates the utility of multi-scale module kernel in discovering disease genes, which could provide insights for the research of relevant issues.

## 1 Introduction

Biomedical data mining plays a vital role in advancing biomedical research, particularly with the rapid growth of biomedical data such as protein-protein interactions (PPI) ^[1]^, disease-related genes ^[2]^, and disease-related phenotypes/symptoms ^[3, 4]^, which has driven the evolution of network medicine ^[5-7]^. To uncover hidden knowledge within these interconnected datasets, numerous network-based mining strategies have been developed. Among these, identifying disease-related genes stands out as one of the most significant areas of focus in the field ^[8-11]^.

The occurrence and development of a lot of diseases are caused by the dysfunctions of genes. Genes as well as their products typically achieve biological functions by synergistic interactions, forming complex biological molecular networks. The dysfunctions of some genes can lead the local perturbation in biomolecular networks, resulting in the phenotypes or symptoms of certain disease ^[12]^. Therefore, a lot of network-based methods have been presented to infer disease-related genes ^[10, 13, 14]^. Generally, there are more interactions or correlations among genes related to the same disease, because they may have more similar functions ^[15]^. Both direct interaction and network distance or closeness are natural strategies for inferring potential disease-related genes ^[16, 17]^, while network propagation is applied more widely to disease-gene identification and related research ^[13, 14, 16, 18-21]^, since it can utilize the information of neighborhood around seeds in biomolecule networks more effectively. However, because of data noise in biomolecule networks, these methods still have room for improvement, which could be achieved through approaches like network enhancement ^[9, 22]^.

The modular structure is one of common properties of biological networks, which can offer valuable insights for the research of complex diseases ^[23, 24]^. Some studies have analyzed network modules for diseases ^[23, 25, 26]^, but the task of mining network modules remains an open challenge ^[27-30]^. For example, we found that the intrinsic resolution limitations may exist in many algorithms, leading to some unexpected results ^[31-33]^. Certain algorithms may produce large-scale network modules that include numerous genes unrelated to the disease. Conversely, some algorithms might generate numerous fragmented network modules, potentially causing the missing of candidate genes that could be relevant to this disease. Moreover, many complex diseases often are related to several modules, as the associated genes may have diverse functions and contribute to the disease through various mechanisms ^[34]^. These modules, while not entirely independent, may be related at a higher functional level or scale. Modules at different scales can offer valuable insights at various levels of analysis. Thus, a multi-scale approach to module analysis could provide more insights for studying these diseases.

Biomolecular networks commonly have multi-scale module structures, consisting of sub-networks that are to recursively split into smaller ones, forming a kind of hierarchical structure ^[35]^. For example, some complexes are composed of several sub-complexes, while function annotations of genes and proteins also have hierarchical organization, suggesting the presence of functional modules across various scales ^[36-38]^. Some studies have shown the significance of these multi-scale modules in understanding biological processes. For instance, Lewis et al. explored the relationship between multiscale modules and functions, highlighting how different scales of modules are linked to specific proteins and processes ^[39]^. Similarly, Wang et al. demonstrated that the hierarchical structure of identified modules roughly corresponds to the hierarchy of Gene Ontology (GO) ^[40]^. In summary, multi-scale module structures offer rich information spanning from local to global structures, which can be valuable for disease research. However, effectively utilizing multi-scale module structures for disease-gene identification is still a challenging task.

To utilize the multi-scale module structures to enhance the capability of identifying disease-related genes, a kind of novel hybrid method called HyMSMK is proposed by incorporating multi-scale module kernel (MSMK) in biomolecule networks. The rest of the work is arranged as follows. Section 2 introduces datasets, including the pre-computation methods. Then, our hybrid methods for incorporating MSMK will be introduced, including the mining of multi-scale module profile, the construction of MSMK, and multiple schemes for incorporating MSMK. Section 3 will introduce experimental settings. Subsequently, we investigate the performance of these incorporating schemes, analyze the effect of kernel sparsification on reducing the requirement for space and time, and show the performance of our method compared to other state-of-art network-based algorithms. Additionally, we perform case studies on several diseases, which identify candidate gene sets for these diseases and validate their relevance to diseases. Finally, we present the conclusions of this study.

## 2 Materials and Methods

Here, the datasets used in this study will be introduced, including datasets of disease genes, disease-related symptoms/phenotypes and PPIs, respectively. Then, we introduce the details of our methods: 1) methods estimating disease-disease associations, 2) methods mining multi-scale module profiles, 3) methods constructing MSMK, and 4) methods incorporating MSMK, leading to several variants (**Table 1**).

**Table 1.** Schemes incorporating multi-scale module kernel.

| Scheme | Description |
| --- | --- |
| CPF | Two heterogeneous networks are constructed, one of which includes multi-scale module kernel, and two ranking lists from them are incorporated by rank aggregation. |
| FS | A heterogeneous network with the fusion of GGA and nGGA is applied. |
| MG | A heterogeneous network with a multi-graph of GGA and nGGA is applied. |

**Table 1.**
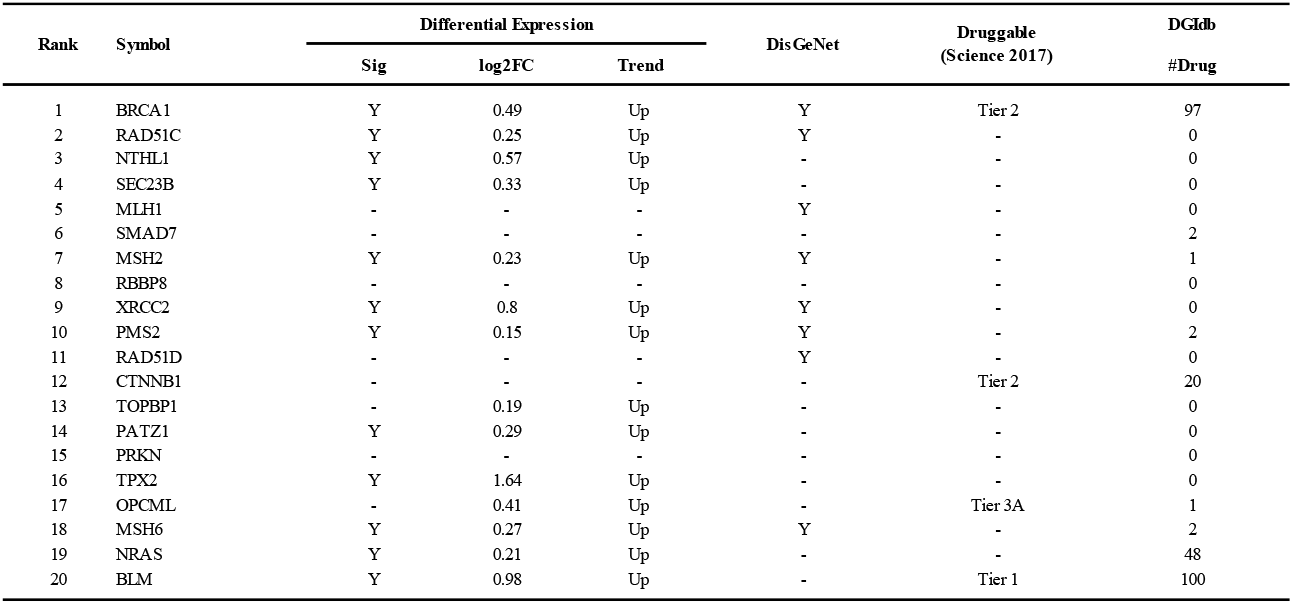
For Breast cancer, candidate genes generated by our method.

| Rank | Symbol | Differential Expression |  |  | DisGeNet | Druggable<br>(Science 2017) | DGldb<br>#Drug |
| --- | --- | --- | --- | --- | --- | --- | --- |
|  |  | Sig | log2FC | Trend |  |  |  |
| 1 | BRCA1 | Y | 0.49 | Up | Y | Tier 2 | 97 |
| 2 | RAD51C | Y | 0.25 | Up | Y | - | 0 |
| 3 | NTHL1 | Y | 0.57 | Up | - | - | 0 |
| 4 | SEC23B | Y | 0.33 | Up | - | - | 0 |
| 5 | MLH1 | - | - | - | Y | - | 0 |
| 6 | SMAD7 | - | - | - | - | - | 2 |
| 7 | MSH2 | Y | 0.23 | Up | Y | - | 1 |
| 8 | RBBP8 | - | - | - | - | - | 0 |
| 9 | XRCC2 | Y | 0.8 | Up | Y | - | 0 |
| 10 | PMS2 | Y | 0.15 | Up | Y | - | 2 |
| 11 | RAD51D | - | - | - | Y | - | 0 |
| 12 | CTNNB1 | - | - | - | - | Tier 2 | 20 |
| 13 | TOPBP1 | - | 0.19 | Up | - | - | 0 |
| 14 | PATZ1 | Y | 0.29 | Up | - | - | 0 |
| 15 | PRKN | - | - | - | - | - | 0 |
| 16 | TPX2 | Y | 1.64 | Up | - | - | 0 |
| 17 | OPCML | - | 0.41 | Up | - | Tier 3A | 1 |
| 18 | MSH6 | Y | 0.27 | Up | Y | - | 2 |
| 19 | NRAS | Y | 0.21 | Up | - | - | 48 |
| 20 | BLM | Y | 0.98 | Up | - | Tier 1 | 100 |

### 2.1 Datasets

In our experimental study, we utilize two datasets of disease-gene associations (DGA). Dataset 1: This integrated dataset combines disease-related genes from OMIM and GWAS ^[41, 42]^. The MeSH system is used to standardize disease nomenclature from multiple sources, and diseases with 20 or more genes are included ^[26, 43]^. Dataset 2: This dataset is derived exclusively from the OMIM database, a comprehensive resource on human diseases and their causal genes, including extensive information on human genes and genetic disorders ^[42]^.

To estimate the associations between diseases, we will extract the data of disease-related symptoms and phenotype abnormalities from human phenotype ontology (HPO) and literature ^[4, 44]^. Complex interactions among proteins form complex protein-protein interaction (PPI) network ^[45, 46]^, which can uncover intrinsic gene-gene associations (GGA) and has become an important basis for the study of molecular mechanisms of diseases. Due to the wide existence of incompleteness of data, we will utilize a comprehensive high-quality interactome constructed by integrating multiple sources of PPIs (e.g., BioGRID, MINT, IntAct, and HPRD) ^[43]^.

### 2.2 Estimation of associations between diseases

#### 2.2.1 Estimation based on co-occurrence between diseases and symptoms

Symptoms are the observable high-level manifestations of a disease and play a critical role in clinical diagnosis and treatment. Research has shown that symptom-based disease similarity is closely linked to molecular interactions ^[4]^. In this study, we are to utilize disease-disease associations inferred by disease-related symptoms from text mining.

Specifically, the associations between diseases and symptoms are characterized by the co-occurrence of MeSH terms for diseases and symptoms, measured by the number of PubMed articles where these terms appear together ^[42]^. A disease is characterized by a vector representation of its associated symptoms, *c*_•,*x*_ = (*c*_1,*x*_, *c*_2,*x*_, …, *c*_*j,x*_, …, *c*_*n,x*_)^*T*^, where *c*_*j,x*_ denotes the co-occurrence between symptom *j* and disease *x*. A co-occurrence matrix of MeSH terms for diseases and symptoms can be constructed by these vectors,

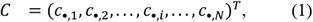

Symptom overlap between diseases can inherently indicate associations between them. However, various symptoms, such as pain and stupor, often vary in their prevalence across diseases, and publication biases may favor certain diseases. To address this heterogeneity, the strength of the association between a disease x and a symptom *j* is typically calculated by the term frequency-inverse document frequency (*TF-IDF*) method ^[47]^. Specifically, a normalized co-occurrence matrix can be obtained by,

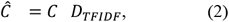

where the diagonal matrix *D*_*TFIDF*_ has diagonal elements:(*D*_*TFIDF*_)_*jj*_ = log(*N*/*N*_*j*_), *N* is the number of diseases and *N*_*j*_ ≥ 1 is the number of diseases having symptom *j*. The similarity scores between diseases can be calculated using the normalized co-occurrence matrix,

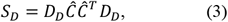

where *D*_*D*_ is a diagonal matrix with diagonal elements: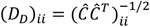. The similarity scores between diseases are between 0 and 1. It is important to note that the above calculation does not consider the inherent correlations between symptoms.

#### 2.2.2 Estimation based on human phenotype ontology

Diseases are generally linked to a group of abnormal phenotypes. The Human Phenotype Ontology (HPO) database offers annotations of human phenotypes for diseases, represented by a collection of HPO terms, which are organized into a directed acyclic graph (DAG) ^[44]^. Unlike the co-occurrence data of diseases/symptoms discussed above, HPO offers more detailed information about the phenotypic abnormalities associated with diseases, offering an alternative approach to uncovering disease-disease associations.

For diseases in OMIM ^[42]^, we assess the disease-disease associations by using dataset from HPO, since it provides phenotype annotations for OMIM disease entries. The similarity scores between diseases can be determined by the number of shared HPO terms. However, not all HPO terms are equally informative. Diseases sharing rare HPO terms are generally more similar than those sharing common ones. To address this, the relevance of HPO terms is quantified using the relative information content (IC) of the terms ^[48]^,

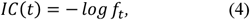

where the frequency of HPO term t is denoted as f_t_. The similarity between two terms t_1_ and t_2_ is derived by,

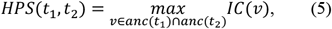

where *anc*(*t*_1_) is the ancestor terms of *t*_1_ on the HPO structure. The similarity score between diseases can be calculated by,

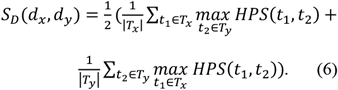

where diseases *d*_*x*_ and *d*_*y*_ are characterized by two sets (*T*_*x*_ and *T*_*y*_) of HPO terms. The similarity scores between diseases are determined by the overall rarity of their shared HPO terms. This measurement of disease-disease similarity incorporates not only the human phenotype annotations but also their inherent relationships within the DAG structure of HPO.

### 2.3 Multi-scale modularity optimization with exponential sampling for mining multi-scale module profile

The mining of multi-scale module structure is an open issue ^[27, 49]^, while it is widely present in biomolecule networks. Modularity optimization (MO) has been widely applied in many fields, including bioinformatics, though there are many network module mining algorithms. Therefore, we will extract modules at different scales by MO, along with exponential sampling (ES) (see **Fig. 1**) ^[50, 51]^. Generally, MO will split a network into module partition by maximizing modularity function Q, that is, finding optimal solution of Q [50-52].

**Fig. 1.**
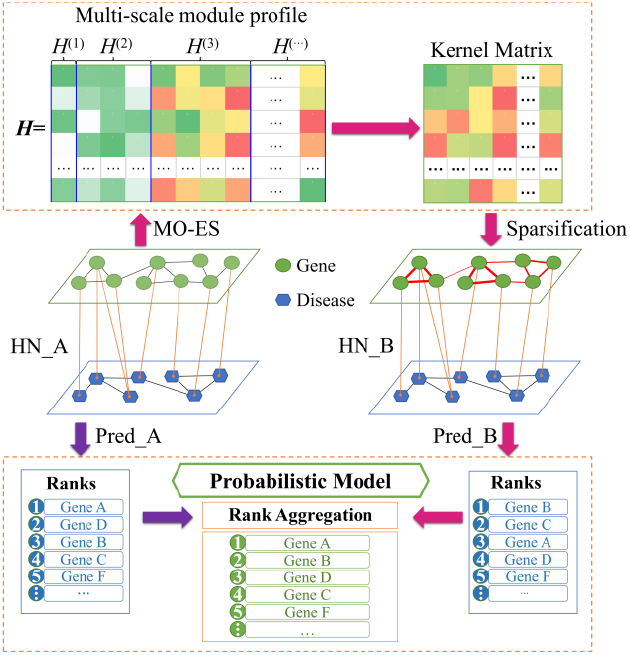
Framework of our method for incorporating multi-scale module kernel by rank aggregation.

For a given module partition of a network, the modularity *Q*, which is widely used as target function of network module mining, can be written as,

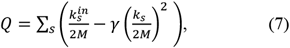

where the total degree of module s is calculated by,

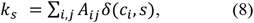

and the total inner degree of module s is calculated by,

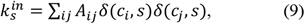

Here, *m* is the number of edges in the network; *A* denotes the adjacent matrix of the network; *c*_*i*_ is the module index of node *i* ; *γ* denotes the resolution parameter. More concisely, it can be written as 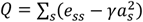, where *e*_*st*_ indicates the fraction of edges between modules *s* and *t*, and *e*_*ss*_ indicates the fraction of ends of edges on module *s*. Given different *γ*-values, one can obtain modules at different scales by feeding a network into effective algorithms for MO ^[49, 53]^. Generally, the smaller *γ* -value will discover larger modules, while the larger *γ*-value will discover smaller modules. A set of single-node modules will be discovered if the *γ*-value is very large, e.g., 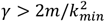, where *k*_*min*_ is the minimum of node degrees ^[54]^. In this study, the Louvain algorithm for MO was used, since it is a fast and efficient algorithm widely used in literature.

To extract a set of meaningful module partitions from local to global scales, according to our previous work, we first define a meaningful range of the values for resolution parameter *γ* ∈ [*γ*_*min*_, *γ*_*max*_], which should be able to include all possible resolution samplings. Given a network, *γ*_*min*_ and *γ*_*max*_ can be denoted as 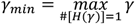 and 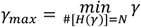, where #[*H*(*γ*)] is the number of modules in the partition *H*(*γ*) and *N* is the number of nodes in the network. Generally, it is difficult to get the accurate boundaries. Therefore, we use semi-empirical boundaries according to previous research ^[55]^. The value of the resolution parameter is in continuous and real resolution space (*γ* ∈ [*γ*_*min*_, *γ*_*max*_]). Continuously scanning all resolution parameter values is meaningless because there must be a lot of information redundancy, and the computer cannot directly process it. To extract meaningful module partitions at different scales, it is necessary to select a series of discrete resolution parameter values for appropriate scales and then extract network module partitions at different scales. Here, we apply the ES strategy to extract a group of discrete *γ*-values in the continuous space, because it can sample meaningful scales reasonably, based on our previous studies ^[51, 53]^. The ES strategy will obtain a group of *γ*-values equally spaced on a logarithmic scale, i.e., *log γ*_*h*+1_ − *log γ*_*h*_*=C*. According to the set of sampled *γ*-values, we apply the MO algorithms to extract a set of corresponding module partitions. Consequently, we have formed a framework of multi-scale MO with ES (i.e., MO-ES) for extracting multi-scale modules.

By the MO-ES framework, we can reasonably generate a series of meaningful network partitions, which can cover the information from the local scales to global scales. For convenience of the quantitative description of module partitions extracted at different scales, we provide the following definitions.

**Definition 1**. We define the following module partition matrix for scale *h* by,

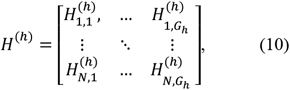

where 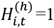 if module *t* in partition *h* contains node i, otherwise 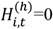.

**Definition 2**. We define the multi-scale module profile by,

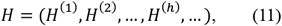

which incorporates multiple module partition matrices at different scales.

Definition 1 can quantitatively characterize module partition at each scale. Definition 2 is used to effectively organize a series of module partitions extracted by MO-ES. It forms the multi-scale module profile, including information from local scales to global scales that may be useful for disease-gene identification. The multi-scale module profile can be viewed as a matrix for multi-scale features, each of which corresponds a column recording the information of the feature.

### 2.4 Construction of multi-scale module kernel

#### 2.4.1 Multi-scale module kernel

The multi-scale module profile can be viewed as a kind of feature matrix containing multi-scale information, which may be useful for uncovering associations between genes. However, like the DSA associations as well as HPA annotations mentioned above, some of these features may not be more informative than others. Genes should be regarded as more similar if they share more infrequent features than others. This is because infrequent features tend to be related to local-scale modules at higher resolutions, usually exhibiting higher edge densities and greater functional homogeneity. Consequently, genes belonging to a module of this kind are likely to be more like each other. To account for this, we use the relative information content (IC) to process these features. A similarity matrix for genes will be constructed by using the processed features. For simplicity in computation, we introduce several definitions.

**Definition 3**. We define a diagonal matrix (*M*_*P*_) of multi-scale module profile by assigning the diagonal element,

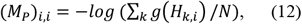

where *g*(·) is defined as a counting function; *g*(*t*) = 1 when *t* > 0, while *g*(*t*) = 0 when *t* ≤ 0.

**Definition 4**. We define a multi-scale module kernel by,

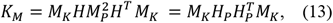

where H_P_ = HM_P_ is a pre-processed multi-scale module profile by the relative information content; *M*_*K*_ denotes a diagonal matrix with the diagonal element,

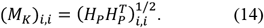

The multi-scale module kernel has the values of elements ranging from 0 to 1, representing a kind of similarity scores between genes. This kernel reveals novel potential associations between genes from the standpoint of multi-scale module structure, offering valuable insights that may aid in identifying disease-associated genes.

#### 2.4.2 Kernel sparsification

The multi-scale module kernel captures potential relationships between genes that are derived from multi-scale module profile. However, it is costly as a full matrix in terms of storage and computation. To address this, we convert it into a sparse matrix by reserving only associations between a gene and its k-nearest neighbors (knn). This kernel sparsification can be defined as follows.

**Definition 5**. We define a ranking vector for the neighbor genes of gene *i*, based on the descending order of the similarity scores between this gene and its neighbor genes, by,

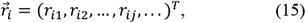

where *r*_*ij*_ denotes the ranking index of neighbor *j* of gene *i*.

**Definition 6**. We define a ranking matrix of genes as the union of the ranking vectors of genes by,

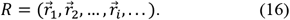

Finally, we generate a new gene-gene association (nGGA) network *G* = (*V, E, W*) based on the full matrix of kernel and its ranking matrix, where *V, E* and *W* respectively indicate the sets of gene nodes, edges and weights,

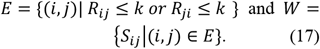

where k denotes the number of the nearest neighbors and *S*_*ij*_ = (*K*_*M*_)_*ij*_ indicates the similarity between nodes *i* and *j*. The nGGA network can be denoted by an adjacent matrix *A*_*M*_ with elements indicating the weights of edges.

The kernel sparsification can be characterized by the number of k-nearest neighbors, the degree of which is inversely related to the number of k-nearest neighbors retained during the sparsification process. This approach helps filter out noise from the data while preserving the essential gene-gene associations identified through multi-scale module profile.

### 2.5 Approaches incorporating multi-scale module kernel

Gene-gene associations provide crucial information for network-based approaches to identifying disease genes. The kernel of multi-scale modules outlined above offers a novel perspective on relationships between genes, derived by using the multi-scale modular structure of the network. This may provide complementary information that aids in the identification of disease genes. In this context, we construct three kinds of hybrid approaches that incorporate the multi-scale module kernel, including rank aggregation model, fusion network model, and multi-graph model (see **Table 1**).

#### 2.5.1 Hybrid approaches based on rank aggregation

The approaches will generate two dual-layer heterogeneous networks of diseases and genes (see **Fig. 1** and **Algorithm 1**). In HN_A, the original gene-gene associations (GGA) are linked to disease-disease associations (DDA) via gene-disease associations (DGA), while in HN_B, nGGAs of multi-scale module kernel are connected to DDAs through DGAs. Next, two lists of prediction genes associated with a disease under study will be generated by conducting the random walk on two heterogeneous networks, separately. Finally, a probabilistic model for rank aggregation is designed to combine two lists of prediction genes. To conduct the random walk on the dual-layer heterogeneous network, we generate a transition matrix based on two intra-layer transition matrices and two inter-layer transition matrices.

##### Algorithm 1 HyMSMK 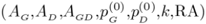

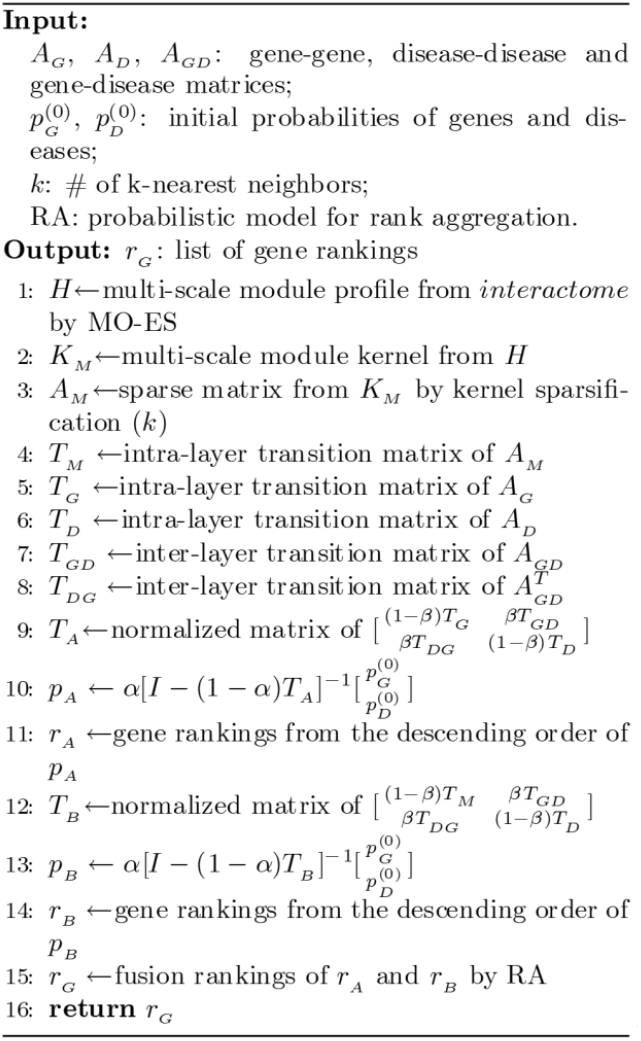

**Definition 7:** We define the intra-layer transition matrix for the layer of gene-gene network as,

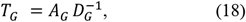

where *A*_*G*_ denotes the adjacent matrix of GGA or nGGA; *D*_*G*_ denotes a diagonal matrix with diagonal elements 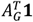 (note that **1** is an all-one vector).

**Definition 8**. We define the intra-layer transition matrix for the layer of disease-disease network as 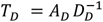, where *A*_*D*_ denotes the adjacent matrix of DDA, and *D*_*D*_ denotes a diagonal matrix with diagonal elements 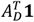.

**Definition 9**. We define the inter-layer transition matrix from diseases to genes as,

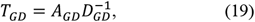

where *A*_*GD*_ denotes the matrix of gene-disease associations: *A*_*GD*_(*j, i*) = 1 when gene *j* is known to be related to disease *i*, and *M*_*GD*_(*j, i*) =0 otherwise; *D*_*GD*_ denotes a diagonal matrix with diagonal elements 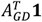.

**Definition 10**. We define the inter-layer transition matrix from genes to diseases as 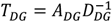, where 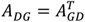 and *D*_*DG*_ denotes a diagonal matrix with diagonal elements 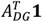.

**Definition 11**. We define the transition matrix of the heterogeneous network as,

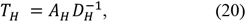

where *D*_*H*_ denotes a diagonal matrix with diagonal elements 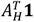, and *A*_*H*_ is the raw transition matrix of the heterogeneous network,

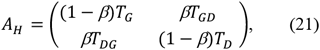

where *β* denotes the probability of jumping between layers.

By the transition matrix *T*_*H*_, one can easily obtain stationary probability vectors of walker arriving at different nodes (genes and diseases). One can calculate its closed solution by,

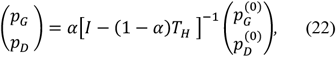

where 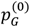 and 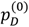 are the initial probability vectors of gene layer and disease layer, respectively; 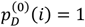 for disease *i* under study, while 0 for other diseases; *α* denotes the restart probability. In order to avoid matrix inversion, one can calculate the stationary probability vectors based on an iterative process,

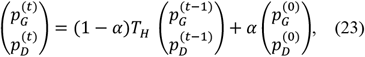

where 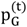 and 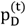 denote the probability vectors of walker arriving at genes and diseases at the t-th time. Based on the descending order of stationary probability, one can obtain a ranking list of genes to identify genes related to the disease under study.

#### Model for rank aggregation

After obtaining two ranking lists of genes (**Fig. 1)**, we generate the final results through a kind of rank aggregation, which is an interesting topic in its own right ^[56]^. Assuming that two lists of genes mentioned above are each obtained based on different features, we can construct a rank-aggregation (RA) model based on Bayes theorem,

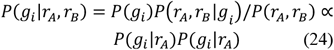

where *P*(*g*_*i*_) denotes the prior probability of gene i being associated with a disease; *P*(*g*_*i*_|*r*_*A*_) denotes the conditional probability of gene i under known ranking *r*_*A*_. To calculate the values of the conditional probability function (CPF), the formal definition of CPF is needed. For each list of prediction genes, a higher ranking of gene often implies a higher probability of being related to the disease. Here, three kinds of CPF’s definitions are designed based on gene rankings:

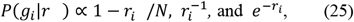

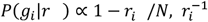, and 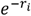, (25) which are denoted respectively by CPF1, CPF2 and CPF3. Here, *r*_*i*_ denotes the ranking of *g*_*i*_. The CPF’s definitions are applied to the rankings *r*_*A*_ and *r*_*B*_ from different heterogeneous networks, respectively. The hybrid approaches with different CPF’s definitions correspond to different variants of our method.

#### 2.5.2 Other approaches incorporating multi-scale module kernel

In addition to the above approaches based on rank aggregation, we have also considered other approaches incorporating multi-scale module kernel, e.g., based on multigraph model and fusion network model.

For approaches based on multi-graph model (MG), a gene-gene multi-graph (MG) need to be generated first [21]. Here, we use GGA and nGGA to construct MG, and generate a multi-graph heterogeneous network by linking MG and DDA with DGA. Then, the list of predictive genes is obtained by the random walk in the multi-graph heterogeneous network.

In order to conduct the random walk on heterogeneous network of multi-graph, the transition matrix T_G_ of GGA is calculated by Eq.(18), and similarly the transition matrix T_M_ of nGGA is calculated by,

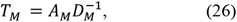

where *A*_*M*_ is the adjacent matrix of nGGA, and *D*_*M*_ is a diagonal matrix with diagonal elements *A*^*T*^ **1**. A transition matrix of the multi-graph is calculated by,

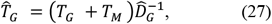

where 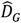 denotes a diagonal matrix and its diagonal element 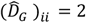 for node *i* located at both the two networks; or 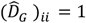 for node *i* isolated in either of networks. By substituting *T*_*G*_ by 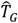 in Eq.(21) and Eq.(20), the transition matrix of the multi-graph heterogeneous network can be constructed. After the iterative process, stationary probability vector of genes can be obtained to identify genes related to the studied disease.

Moreover, we also consider the hybrid approaches based on fusion network model as comparison. We start by constructing a fusion network (FS) that combines edges from both GGA and nGGA. Next, we link the FS network with DDA via DGA to form a dual-layer heterogeneous network. Finally, we apply the previously mentioned random walk to this network to obtain a list of predictive genes associated with the studied disease.

## 3 Experimental Results

This section provides a systematic evaluation of performance for our HyMSMK method, including its three variants using different schemes. First, the experimental settings are introduced. Then, through traditional cross-validation experiments, we analyze the effects of different integration schemes and kernel sparsification, compare our method against state-of-the-art network-based algorithms across multiple disease-gene datasets. Finally, we present multiple case studies that demonstrate the disease relevance of predicted candidate genes.

### 3.1 Experimental settings

In this study, we employ two evaluation schemes, two types of control sets, and three kinds of standard metrics to evaluate the performance of methods.

#### (1) Two evaluation schemes

We employ classical 5-fold cross-validation (5FCV) to evaluate our method. For 5FCV, applied to Dataset1 and Dataset2, the known disease-related genes are randomly divided into five subsets. Each subset is used as the test set once, and the rests are to form the training set. Additionally, we demonstrate the effectiveness of our method through case studies.

#### (2) Two types of control sets

Both a test set and a control set for candidate genes are needed to construct a candidate gene set for evaluation. Two types of control sets will be used, including the artificial linkage-interval control set denoted as ALICS and the whole-genome control set denoted as WGCS. ALICS has 99 nearest genes for each test gene on the same chromosome [16]. All unknown genes are used as the control set in WGCS, excluding those in the training and test sets.

#### (3) Three kinds of standard metrics

After obtaining the predictive list of candidate genes, this study will use three standard metrics (Recall, Precision and AUPRC) to evaluate the performance of predictive algorithms. a) Recall measures the proportion of known disease genes identified within the ranking list of top-k candidate genes, with regard to the test set. b) Precision indicates the likelihood of identified known disease genes within the ranking list of top-k candidate genes. Both Recall and Precision change with the length of the top-k ranking list, allowing for a straightforward comparison of the algorithms’ local performance. c) To assess overall algorithm performance, we calculate the area under the Precision-Recall curve (AUPRC), where the PRC curve plots Precision on the *y*-axis and Recall on the *x*-axis.

### 3.2 Comparison between different schemes incorporating multi-scale module kernel

We have designed different schemes to incorporate multi-scale module kernel, leading to several variants of our method. The predictive performance of the schemes will be compared under different scenarios (**Fig. 2)**. For both control sets (ALICS and WGCS), the CPF2 scheme outperforms all others across the three kinds of evaluation metrics in Dataset1, and it can yield better or comparative results than other schemes in Dataset2 (see Supplemental materials). Therefore, CPF2 is selected as the default scheme for the subsequent studies.

**Fig. 2.**
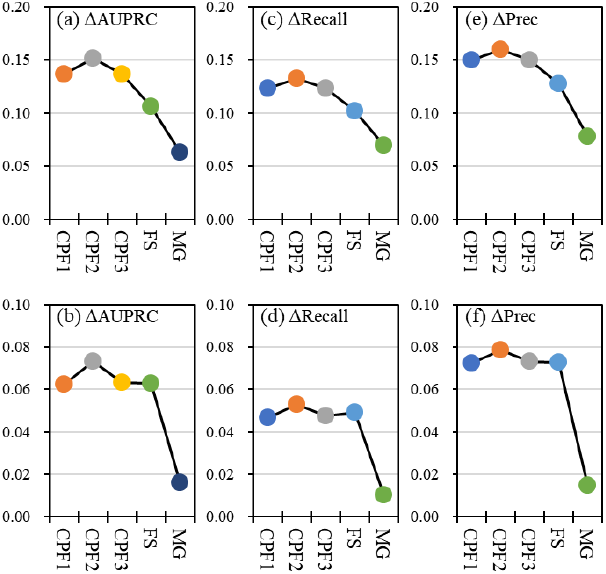
Performance increase of HyMSMK with distinct schemes incorporating multi-scale module kernel, relative to baseline, in Dataset1. (a)-(b) ΔAUPRC, (c)-(d) ΔRecall, (e)-(f) ΔPrec. (a), (c) and correspond to the ALICS control set, while (b), (d) and f) correspond to the WGCS control set.

Additionally, the schemes based on k aggregation generally perform better than MG and FS. This may be because FS and MG can introduce significant interference when fusing the original GGA and nGGA networks. The MG scheme will switch the random walkers between networks with equal probability, and the predictive scores from each network are to be integrated immediately after each step. The FS scheme directly combines the GGA and nGGA networks into one and results in the loss of unique information for each. By contrast, the CPF schemes treat each network as an independent individual, minimizing interference between them.

### 3.3 Effect of kernel sparsification

The kernel sparsification process is helpful for improving the storage and computing of multiscale module kernel and filtering out data noises. In this section, we analyze the effect of the kernel sparsification process.

The results show that there are clear and consistent upward trends under both control sets (**Fig. 3**). The decrease of sparseness, that is, the increase of k-nearest neighbors (knn), will lead to the increasing performance of our method. It is more susceptible to small knn-values. A possible reason is that knn increasing from a low value rapidly introduces more strongly correlated information, leading to a faster improvement in performance. However, for very large knn-values, it can only add some neighbors with low similarity scores to each gene, causing the performance improvement to level off. Based on the performance trends, we can see that a reasonable choice of knn is 1000 in this study, striking a good balance between performance and speed, as it represents a turning point in the performance curve. Moreover, the performance of our method also improves rapidly as knn increases and stabilizes at a high level in Dataset2 (see Supplemental materials). Setting knn11000 proves to also be an effective choice for our method, despite a slight decrease in Recall at this point. Notably, the AUPRC continues to improve as knn increases. It is important to mention that our method with knn10 is equivalent to the reference method, indicating that our method with knn11000 significantly outperforms the reference method.

**Fig. 3.**
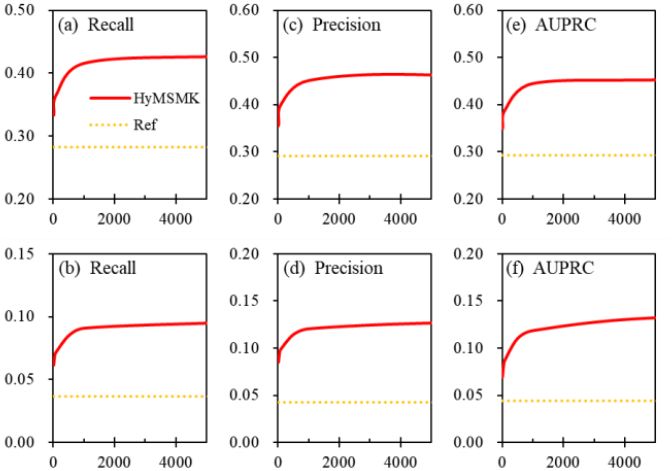
Effect of kernel sparsification on predictive performance: (a)-(b) Recall, (c)-(d) Prec, (e)-(f) AUPRC (Dataset1). (a), (c) and (e) correspond to the ALICS control set, while (b), (d) and (f) correspond to the WGCS control set.

Here, the requirements of space and time for our method under different levels of kernel sparsity will be evaluated, using both sparse matrix representation mode (sMRM) and full matrix representation mode (fMRM). We can see that usage of memory in sMRM increases approximately linearly with knn, and time consumption follows a similar trend, growing alongside space requirements (**Fig. 4)**. Notably, when knn is small (e.g., knn≤1000), the memory usage of sMRM is significantly lower than that of fMRM. For example, for the default value of knn11000 derived in the above analysis of performance, the memory usage of MSMK after the sparsification process is about 0.24GB under sMRM, while it is about 1.34 GB under fMRM. As a result, the memory usage of HyMSMK (estimated) and its time consumption are significantly reduced under sMRM, much less than that in fMRM.

**Fig. 4.**
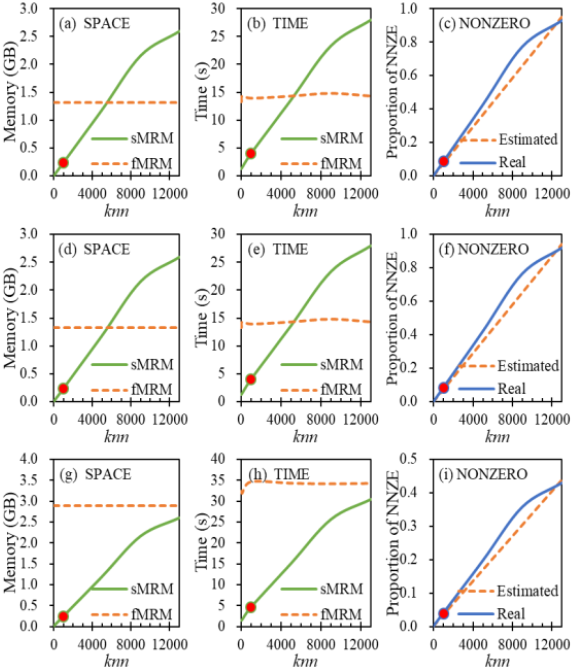
Space and time requirements of HyMSMK under the sparse matrix represention mode (sMRM) and full matrix represention mode (fMRM), as well as the real and estimated proportion of non-zero elements (pNNZE), as a function of knn. (a)-(c) Test in homogeneous network nGGA; (d)-(f) Test in Dataset1; (g)-(i) Test in Dataset2.

We note that the memory usage of HyMSMK using sparsified kernel under sMRM is similar to that under fMRM when knn is close to 5000, and it will become larger with the increase of knn (see **Fig. 4**d-e). This is because the sparsity of the kernel matrix will decrease with the increase of knn, gradually becoming non-sparse, and the forced sparse representation of non-sparse matrix will lead to the increase of memory usage and time consumption, due to the inherent mechanism of sMRM. This is also the case in Dataset2 (**Fig. 4**g-h), but the critical point of knn (where the memory/time usage of sMRM is close to that of fMRM) is delayed. As we see, the memory/time usage of sMRM is still less than that of fMRM when knn is close to the number of nodes in gene network. This is because the size of DDA in Dataset2 is much larger than that in Dataset1, and its DDA and DGA are sparse. In addition, we can see that the growth rate of space consumption slows down when knn is very large, and this is also the case for the growth rate of time consumption. This is because the space and time consumptions are closely related to the number of non-zero elements (NNZE) in matrix. For example, **Fig. 4**c shows that the growth rate of NNZE already slows down when knn is very large, while the growth rate for small value of knn is quicker than the estimates using knn as mean degree of nodes.

In summary, the kernel sparsification process effectively reduces memory usage and time consumption, enhancing its storage efficiency and computational speed. Moreover, HyMSMK utilizing sparsified kernel (e.g., knn11000) maintains stable and high performance in disease-gene identification.

### 3.4 Comparison to other algorithms

We compare the performance of HyMSMK to network-based baselines including RWRH ^[57]^, CIPHER ^[58]^, BiRW ^[59]^, KATZ ^[60]^ and HyMM^[55]^, by the 5FCV experiments. **Fig. 5** demonstrates that our method outperforms the basic baseline algorithms (RWRH, CIPHER, BiRW and KATZ) under ALICS and WGCS control sets. In details, HyMSMK improves the AUPRC-, Recall-, and Prec-values by 41%, 47%, and 37%, respectively, under ALICS, and by 27%, 7%, and 10%, respectively, under WGCS, when compared to the best of the basic baselines. Further, we compare HyMSMK to another multiscale module-based algorithm called HyMM. The results show that HyMSMK outperforms HyMM under ALICS control sets, and they have similar values of AUPRC under WGCS control sets (see **Fig. 5**). The Recall and Precision curves further confirm the strong local performance of our method (**Fig. 6**).

**Fig. 5.**
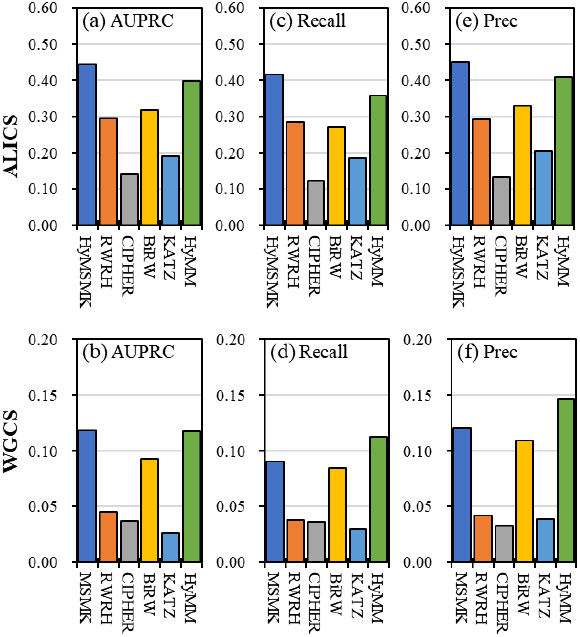
For Dataset1, performance comparison to other algorithms. (a)-(b) AUPRC, (c)-(d) Recall, and (e)-(f) Precision. The first and second rows correspond to the ALICS and WGCS control sets, respectively.

**Fig. 6.**
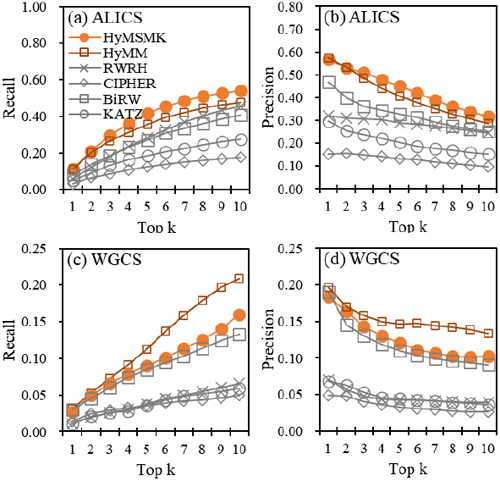
For Dataset1, comparison of local performance (Recall and Precision) of HyMSMK to other algorithms. (a)-(b) ALICS, (c)-(d) WGCS.

The results in **Fig. 7** show that HyMSMK also surpasses all the comparison algorithms under both control sets, in Dataset2. HyMSMK improves AUPRC, Recall, and Precision by 5%, 7%, and 9% under ALICS, and by 27%, 47%, and 48% under WGCS, respectively, when compared to the best of the comparison algorithms. In this dataset, our HyMSMK in terms of AUPRC, Recall, and Precision has consistently better performance than the multiscale module-based algorithm HyMM under both control sets. **Fig. 8** further corroborates HyMSMK’s excellent local performance. Overall, HyMSMK proves to be a consistently strong choice under both control sets.

**Fig. 7.**
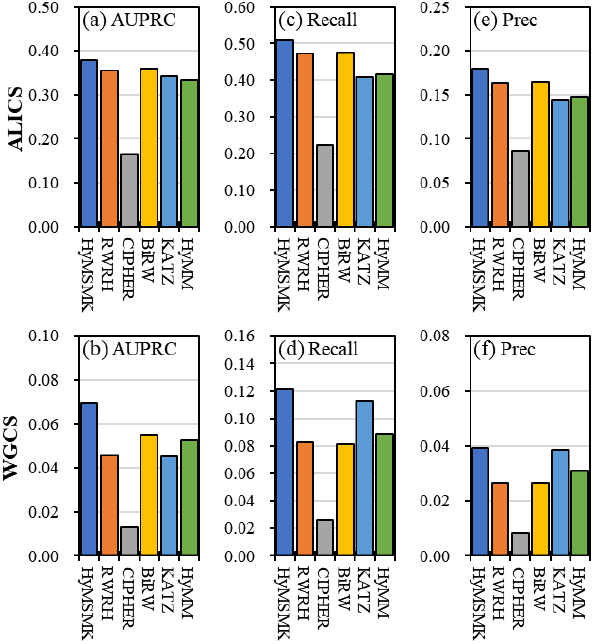
For Dataset2, performance comparison to other algorithms. (a)-(b) AUPRC, (c)-(d) Recall, (e)-Prec. The first and second rows correspond to the ALICS and WGCS control sets, respectively.

**Fig. 8.**
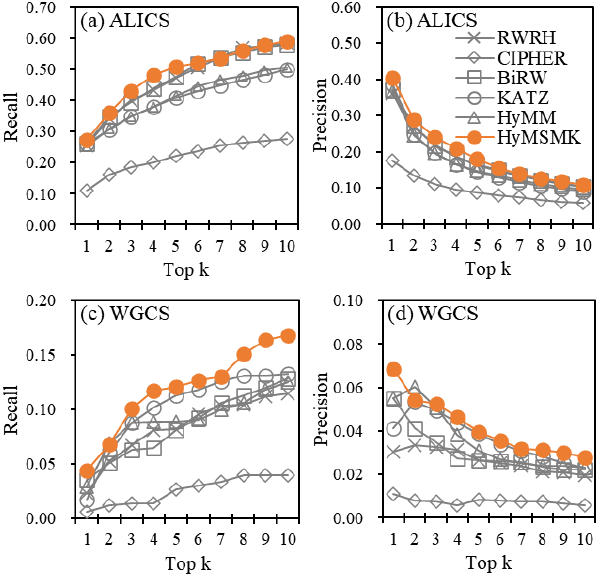
For Dataset2, comparison of local performance (Recall and Precision). (a)-(b) ALICS, (c)-(d) WGCS.

### 3.5 Case studies

We perform case studies for several diseases to demonstrate the effectiveness of our method further. For each disease, a list of candidate genes is generated using the OMIM database as the training set. The disease relevance of these predicted genes is then evaluated through literature validation and differential expression analysis.

**Table 2** presents the predicted top-k candidate genes for Alzheimer’s disease (AD, #104300). Among these, 13 genes have supporting literature records related to AD, including 5 genes with AD-related entries in the DisGeNet database (APOE, TF, PSEN1, MTHFR, and MAOB). Additionally, 7 genes are supported by AD-related literature but are not listed in DisGeNet (EIF4G1, SNCA, SNCB, MAOA, MTHFD1, CP, and AOC3). For instance, the APOE ε2 allele is one of the genetic protective factors in AD ^[61, 62]^; Sutovsky et al showed that the APOE4 homozygotes and the APOE4 carriers with TT genotype of MTHFR rs1801133 have the highest risk of AD ^[63]^. Jouini et al observed that the plasma concentrations of transferrin (TF), ceruloplasmin (CP), and iron in AD group were significantly lower than that of adults with normal cognition, and there was a strong correlation between most of them^[64]^. Zhang et al ^[65]^ found that a mutation (S169del) of PSEN1 can affect the APP processing and Aβ generation, and the neuritic plaque formation and the deficits of learning and memory can be promoted in AD model mice. This suggests that this mutation (S169del) may be an important site of AD treatment as a potential target. Florentinus-Mefailoski demonstrated that an increase in the mean precursor strength of peptides from various plasma proteins, including EIF4G1, is associated with AD ^[66]^. α-Synuclein and tau aggregations are the neuropathological molecular feature of Parkinson’s disease and Alzheimer’s disease, respectively, but they also may co-occur in patients. Bassil showed that the accumulation and spread of tau in mice were weakened due to the lack of endogenous α-Synuclein, but the seeding or spreading ability of α-syn was not affected by the lack of tau. This indicates the important role of α-Synuclein in modulating tau pathological burden and spreads in the brains of patients ^[67]^. Bergström reported the increased levels of synuclein beta (SNCB) in the cerebrospinal fluid (CSF) of AD patients compared to controls ^[68]^. Similarly, Quartey et al. discovered that MAO-A activity increased in the cortical regions of AD patients but not in hippocampal samples, whereas MAO-B activity showed an increase in both regions ^[69]^. Jiang et al showed that the polymorphisms of Methylenetetrahydrofolate reductase (MTHFR) gene: C677T and A1298C were related to AD in the Chinese population ^[70]^; and Zuin et al showed that MTHFR C677T polymorphism had the association trend with late-onset Alzheimer’s disease in Italian population ^[71]^. Bi et al showed that the G1958A A allele of MTHFD1 may be a weak risk factor of early onset AD ^[72]^.

**Table 2.**
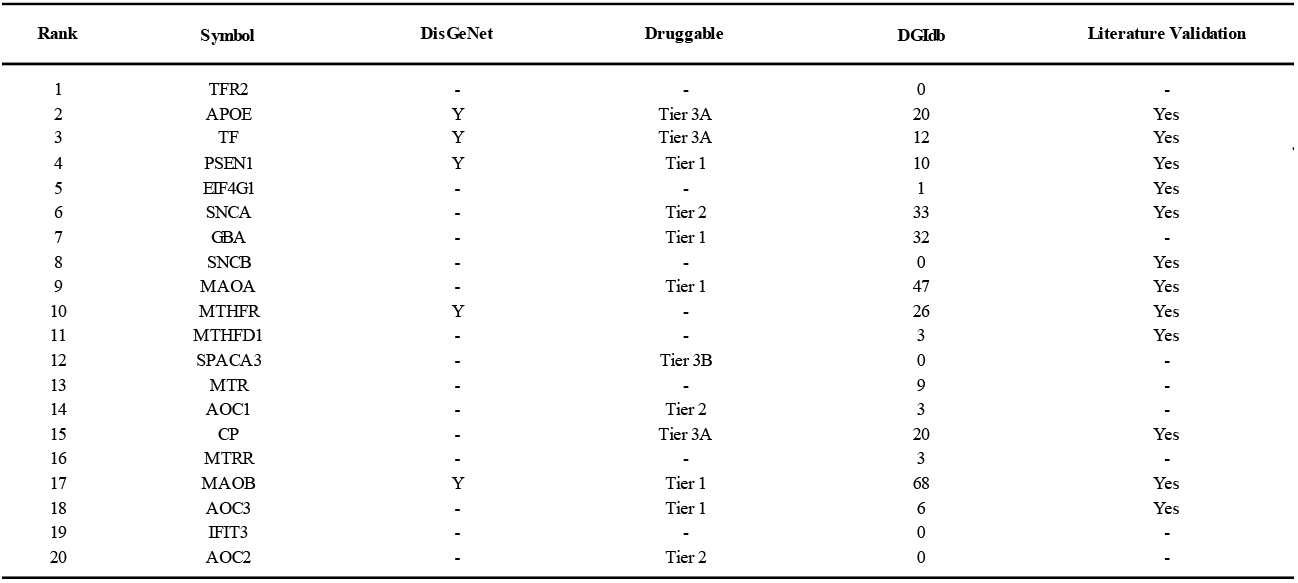
For Alzheimer’s disease, candidate genes generated by our method.

**Table 2.** For Lung cancer, candidate genes generated by our method.

| Rank | Symbol | Differential Expression |  |  | DisGeNet | Druggable<br>(Science 2017) | DGldb<br>#Drug |
| --- | --- | --- | --- | --- | --- | --- | --- |
|  |  | Sig | log2FC | Trend |  |  |  |
| 1 | TP53 | - | - | - | Y | Tier 1 | 383 |
| 2 | DICER1 | - | - | - | - | - | 0 |
| 3 | RB1 | - | - | - | Y | Tier 2 | 23 |
| 4 | BAP1 | - | - | - | Y | - | 6 |
| 5 | TSC1 | - | -0.24 | Down | - | - | 3 |
| 6 | TSC2 | - | - | - | - | - | 1 |
| 7 | MLH1 | Y | -0.3 | Down | Y | - | 0 |
| 8 | POU2F1 | Y | 0.45 | Up | - | - | 0 |
| 9 | MSH2 | Y | 1.13 | Up | - | - | 1 |
| 10 | MUC5B | - | - | - | - | Tier 3B | 0 |
| 11 | SFTPA2 | Y | -3.18 | Down | - | Tier 3B | 0 |
| 12 | HRAS | Y | 0.97 | Up | Y | Tier 1 | 38 |
| 13 | FAS | - | - | - | - | Tier 3A | 19 |
| 14 | NRAS | Y | 0.38 | Up | Y | - | 48 |
| 15 | CHEK2 | Y | 1.87 | Up | - | Tier 1 | 29 |
| 16 | SHOC2 | Y | -0.35 | Down | - | - | 0 |
| 17 | RASGRP2 | Y | -2.5 | Down | - | - | 0 |
| 18 | RAP1GDS1 | - | - | - | - | - | 0 |
| 19 | ERCC5 | Y | -0.33 | Down | - | Tier 2 | 1 |
| 20 | AKT1 | - | - | - | Y | Tier 1 | 78 |

**Table 3.**
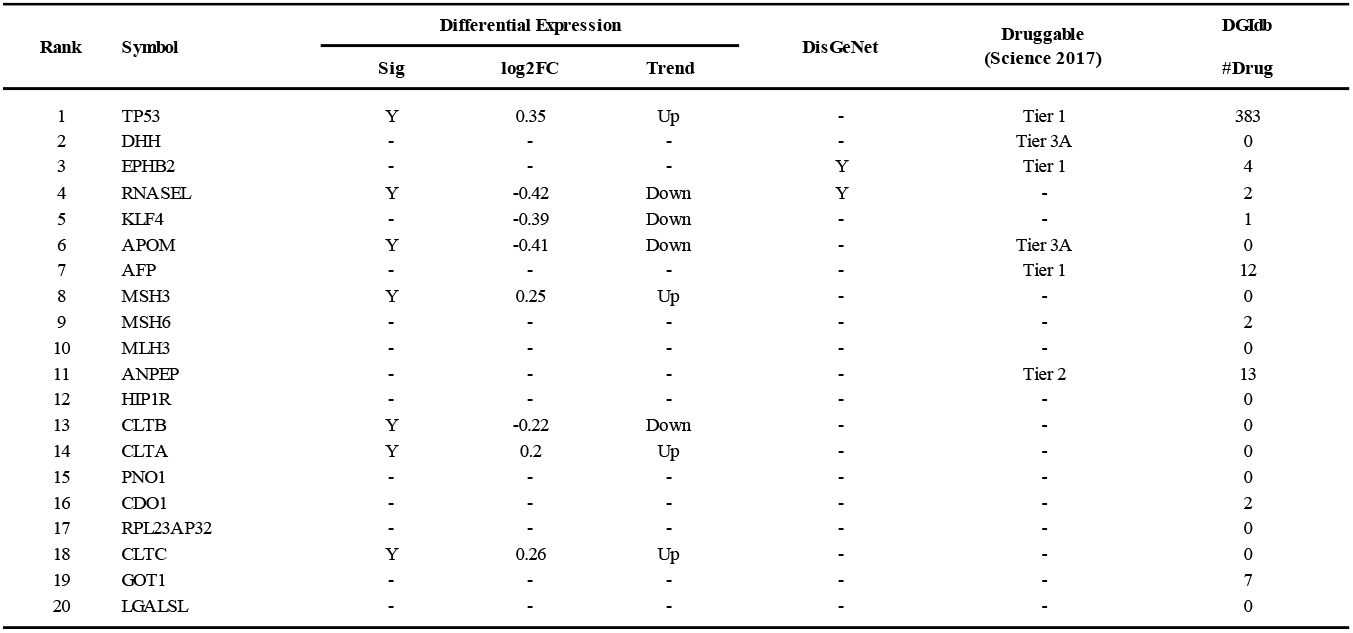
For prostate cancer, candidate genes generated by our method.

Stroke and AD have evident interrelationships mainly because the vascular system dysfunction (VSD) and their pathologies have important effects on cerebral vessels, together with increased SSAO levels. The VSD can directly lead to the occurrence and progression of AD, and there are signs of dysfunction of blood-brain barrier (BBB) in the early stages of AD. Solé et al showed that the expression of SSAO/VAP-1 was related to endothelial activation through changing the release of pro-inflammatory and pro-angiogenic neuroproteins, and revealed the important role of vascular SSAO/VAP-1 in the BBB dysfunction related to AD progression, which provides new insights to the study of AD’s therapeutic targets ^[73]^.

Furthermore, we analyze the top-k candidate genes predicted for other diseases, evaluating their associations with specific conditions through gene differential expression analysis ^[74]^ and the DisGeNet database [2]. shows the predicted top-20 candidate genes for breast cancer (#114480), of which 12 are significantly differentially expressed in breast cancer, and 8 have breast cancer-related entries in the DisGeNet database, suggesting a strong likelihood of their associations with the disease. **Table 2** presents the predicted top-20 candidate genes for lung cancer (#211980), with 10 genes significantly differentially expressed and 7 recorded in the DisGeNet database as associated with lung cancer, indicating their potential relevance to the disease. **Table 5** lists the predicted top-20 candidate genes for prostate cancer (#176807), among which 7 are significantly differentially expressed in prostate cancer, and 2 genes have prostate cancer-related entries in the DisGeNet database, implying a probable link to the disease.

## 4 Conclusion and discussion

Disease-gene discovery is a critical area of biomedical data mining, driven by the high cost and time required for biomedical experiments. The modular structure of biological networks offers valuable information for disease research, which is embedded in multi-scale module structures. However, effectively leveraging the information from multi-scale module structures to identify disease-associated genes remains a challenge. To address this, we have proposed the HyMSMK method of incorporating the multi-scale module kernel. It extracts the multi-scale modules at different scales from a comprehensive interactome through multi-scale modularity optimization and exponential sampling, and then constructs the multi-scale module profile, which can capture multi-scale information from local structure to global ones. The multi-scale module kernel is constructed using the multi-scale module profile, combined with the relative information content (IC), which is further preprocessed by kernel sparsification. Multiple schemes for incorporating multi-scale module kernel are designed to explore more effective approaches for discovering potential disease-related genes.

Through a series of experiments, we have evaluated our HyMSMK method under both ALICS and WGCS control sets systematically and examined the effects of various incorporating schemes and kernel sparsification. The results across various scenarios demonstrated the strong performance of our method, affirming the utility of multi-scale module structure in disease-gene identification. Among the schemes, CPF2 is recommended as it consistently delivers reliable results across various scenarios. Additionally, case studies for specific diseases validate the relevance of candidate genes identified by our method, through literature support and differential expression analysis, further emphasizing our method’s effectiveness.

Multiscale module mining is still an open issue in complex networks, including biological networks. It is based on the given network that the mining of multi-scale modules are conducted. So, the extraction and optimization of multi-scale modules would inevitably be affected by the scale of the data or the sparsity of the network. Our previous theoretical research for module mining algorithms found that existing algorithms based on global target function optimization have two types of resolution limitations. The first type of limitation is that when the network size is very large, some small network modules may be hidden and cannot be directly discovered by the algorithms. The second type of limitation is that although it is possible to identify modules at different scales through algorithms with adjustable resolution, when identifying small-scale modules by increasing resolution, some large modules may experience cracking issues. That is to say, under certain conditions, it is impossible to identify both large-sized and small-sized modules simultaneously. There is still a lot of research to this day in this field of module mining. The analysis and mining of biological networks from the perspective of modules can alleviate the problem that the study of a single node or edge is more easily affected by network noise. Furthermore, compared to traditional hard partitioning methods of network modules, multi-scale module mining can extract multi-scale information from local to global levels. The impact of network sparsity will further be reduced due to its more detailed network mining.

The method in this work has been proved to be able to achieve good performance, while there is still room for further improvement. To utilize the information in multi-scale module structure, we provide an effective strategy in this article, but it does not mean that the information has been fully and effectively utilized. We use the extracted multi-scale information to construct a multi-scale module kernel for disease-gene prediction, but it may also serve as a feature input for machine learning algorithms such as support vector machines, random forests, and neural networks to train machine learning models.

Module structures widely exist in various biological networks, such as PPI networks, metabolic networks, and disease-disease networks. In this work, we only consider the multi-scale modules in GGA networks, while disease-gene prediction may benefit from the mining of multi-scale modules in DDA networks. In disease networks, similar diseases may aggregate into network modules, and there are also varying degrees of similarity between different diseases. From the perspective of different degrees of similarity, disease modules at different levels can be extracted. These pieces of information have the potential to improve disease networks and provide more valuable information for disease-gene prediction. The HyMSMK method uses MO-ES to extract multiscale modules due to its stable performance in our previous research, while it is possible to enhance disease-gene prediction based on multiscale modules by developing more effective module identification algorithms. For instance, Hu et al proposed a method for clustering analysis by exploiting a variety of network motifs which are a kind of small patterns recurring in a network and showed that taking account of higher-order structures could obtain new insight into the analysis of biological networks ^[75]^.

In this work, we only use data for disease-related genes/phenotypes/symptoms and gene-gene networks. In past decades, a large amount of omics data from genomics to transcriptomics and proteomics has been generated continuously ^[76]^. The occurrence of diseases can be reflected at different levels of the organism, from genes to protein complexes and pathways. In theory, the utilization of more relevant omics data, e.g., gene mutations, methylations and transcription factors, may improve the mining of biological knowledge. Disease-related research will benefit from the accumulation in the amount and types of omics data, enhancing the ability of disease-gene prediction^[77]^.

Personalized study of diagnosis and treatment is an inevitable trend in the future development of precision medicine. The availability of patient-level datasets containing both genotypic and phenotypic data offers new opportunities to explore individual pathogenic genes, though most research currently focuses on disease classes or their subcategories. With advancements in single-cell sequencing technologies, vast amounts of cell-level multi-omics data are being generated at an unprecedented rate ^[76, 78]^. Integrating cell-level multi-omics data across various molecular levels remains challenging due to its inherent heterogeneity, while it has the potential to uncover the complexity of biological systems.

In conclusion, this study presents an efficient framework HyMSMK by incorporating the multi-scale module kernel into disease-gene identification, which may offer valuable insights for leveraging multi-scale module structures in the exploration of other bio-entity associations^[79-82]^. It can theoretically be generalized to explore associations between other biological entities, such as disease-metabolite associations, disease-ncRNA associations and disease-drug associations, since the networks consisting of these bio-entities also have module structures.

## Acknowledgments

This work was supported by the National Natural Science Foundation of China under (Grant No.62472051), the Hunan Provincial Natural Science Foundation of China (Grant No. 2024JJ5006), the Research project of Changsha University of Science and Technology (097_000303919), the National Key Research and Development Program of China (Grant No. 2019YFA0706202), the National Natural Science Foundation of China under (Grant No.62202502) and the High Performance Computing Center of Central South University..

## Notes

### Competing Interest Statement

The authors have declared no competing interest.

### Summary of Updates

The authors and affiliations were updated. The manuscript was rewritten.

